# Association of the *GIPR* Glu354Gln (rs1800437) polymorphism with hypertension in a brazilian population

**DOI:** 10.1101/340539

**Authors:** Alexandro Marcio da Silva Mattos, Sarah Conchon Costa, Giovana Outuki, Gustavo Kendy Camargo Koga, Cláudia Nascimento Montemor, Giana Zarbato Longo, Maria Angelica Ehara Watanabe, Marla Karine Amarante, Tânia Longo Mazzuco

**Affiliations:** Post-graduation Program of Health Sciences, Londrina State University, Londrina, Paraná, Brazil.; Endocrine Interactions Research Group, Londrina State University, Londrina, Paraná, Brazil.; School of Medicine, Pontifical Catholic University, Londrina, Paraná, Brazil.; Department of Nutrition and Health, Federal University of Viçosa, Vçosa, Minas Gerais, Brazil.; Laboratory of DNA Polymorphisms and Immunology, Department of Pathological Sciences, Biological Sciences Center, Londrina State University, Londrina, Paraná, Brazil; Division of Endocrinology of Medical Clinical Department, University Hospital, UEL, Londrina, Brazil

## Abstract

**Objective:** To know the prevalence of the Glu354Gln polymorphism of the *GIPR* gene, investigate possible associations with arterial hypertension and relationships with cardiometabolic diseases.

**Method:** A total of 311 subjects recruited from the Clinical Hospital of Londrina State University, located in a Brazilian metropolitan area. Random stratification was performed considering gender and geographic regions. Data were collected through interviews including anthropometric, sociodemographic and metabolic diseases related diseases. In order to analyze *GIPR* Glu354Gln gene polymorphism, polymerase chain reaction followed by followed by restriction fragment length polymorphism (PCR-RFLP) was performed.

**Results:** The highest prevalence for the allele C carriers were found in the Caucasian 29.4% (p = 0.043, OR = 1,89), hypertensive 37.1% (p < 0.0001), smokers 38.3% (p = 0.014) and dyslipidemic group 41.2% (p = 0.019). In this work 46.9% of the participants (n = 146) presented diseases related to cardiometabolic diseases. The results indicated that 60% of hypertensive patients (p = 0.004) and 64.7% of dyslipidemic patients (p = 0.046) were male. Among participants who presented cardiometabolic diseases, arterial hypertension was the most prevalent disease (71.9%), followed by obesity (43.8%). The family comorbidities history to cardiometabolic diseases (DM2, AH, dyslipidemia and obesity) had no significant association with the *GIPR* Glu354Gln genetic polymorphism. Although there was no difference in the case-control analyses for *GIPR* Glu354Gln for cardiometabolic group, regarding C allele carriers there were twice associated with arterial hypertension (p<0,001) and dyslipidemia (p<0,03).

**Conclusion:** The prevalence of the *GIPR* Glu354Gln for the CC genotype and for the C polymorphic allele was 25.7% and 3.2%, respectively. This study shows the potential participation of the *GIPR* Glu354Gln polymorphism with the pathophysiology of arterial hypertension, dyslipidemia in this Brazilian population. Taking into account the rarity of the CC genotype, additional studies with larger numbers of participants could contribute to a better understanding.

## Introduction

Glucose-dependent insulinotropic polypeptide (GIP) is a hormone secreted by K cells from duodenum and proximal jejunum, after glucose and fat uptake. GIP accounts for 80% of β-cell enteric stimulation by increasing pancreatic insulin secretion induced by glucose intake, which is known as the incretin effect (1). Functional GIP receptors play a major role in insulin resistance and hepatic steatosis in high fat feeding mice (2, 3). GIP receptors *(GIPR)* are mainly expressed in islet cells, in addition to osteoblasts, adipocytes, pancreas, lung, adrenal, kidney and thyroid (1). Nonpancreatic *GIPR* function is related to adrenal cell proliferation, GIP-dependent oversecretion of cortisol and aldosterone (4, 5, 6) and neurogenesis in the central nervous system (6).

Several GIP receptor polymorphisms have been described. These genetic variations could play a role on gluco-insulin homeostasis and body composition. Some single nucleotide polymorphisms (SNPs) located in coding regions of the *GIPR* gene (rs8111428, rs2302382 and rs1800437) were related to obesity (7). The *GIPR* rs10423928 was associated with high postprandial glucose and insulin levels (8), decreased lean mass (9) and low BMD in early postmenopausal women (10). The *GIPR* SNP rs1800437 (Glu354Gln) showed a borderline association with cardiovascular disease (CVD) in a study of 200 patients from a Dutch population (11). Genetic variations of the *GIPR* were not associated with childhood obesity; nevertheless, a potential role in glucose homeostasis was described for the *GIPR* SNP Glu354Gln (12).

Our preliminary results involving cardiometabolic risk factors has pointed to the potential role for the *GIPR* Glu354Gln genetic variant in systemic arterial hypertension (13). The aim of the present study was to assess the prevalence of the *GIPR* Glu354Gln in a Brazilian population and to investigate its association with socio-demographic and clinical characteristics related to the cardiometabolic syndrome.

## Material and methods

### Human subjects and blood samples collection

For the study of *GIPR* Glu354Gln gene polymorphism, individuals were recruited from outpatient clinic at the Clinicals Hospital, Londrina State University (UEL). The population consisted in a total of 311 unrelated subjects (151 male) from a Brazilian metropolitan area, with a population of approximately 1.067,214 people. Sample was determined considering a 6% polymorphism prevalence, with a 95% significance level and a 3% precision level; to the results, were added 30% to compensate possible losses. The sampling process was random, 25 male and 25 female subjects for each geographic region of Londrina-PR, previously established (north, south, east, west, center and periphery), seeking criteria that would serve a more reliable representation of the population of this region. It was considered to be a smoker every individual who has used a minimum of 100 cigarettes and still uses them daily or eventually (14). All procedures in the present study were approved by Londrina State University Ethics Committee for Research Involving Human Subjects (CAAEA 46541215.0.0000.5231) and all individuals were informed about the research and freely signed a consent term prior to biological material collection.

### DNA extraction

Genomic DNA was obtained from blood samples using Biopur Mini Spin kit (Biometrix Diagnostica^®^, Curitiba, PR, Brazil). DNA samples were quantified using a NanoDrop2000c Spectrophotometer (ThermoFisher Scientifc, Wilmington, DE, EUA) at 260 nm. The 260/280 nm and 260/230 nm absorbance ratios were measured to assess protein and organic compound contamination, respectively.

#### Genotyping assay

Polymorphism analysis for *GIPR* Glu354Gln was performed by polymerase chain reaction followed by restriction fragment length polymorphism (PCR-RFLP). PCR conduced in a final volume of 25 μL containing 100 ng of DNA template, 1.25U of Taq DNA polymerase, 10% of PCR Buffer (200 mM Tris–HCl pH 8.4, 500 mM KCl), 0,1 mM dNTP, MgCl2 50 mM. All PCR reagents were purchased from Invitrogen (Invitrogen TM, Life Technologies Carlsbad, CA, USA). The following oligonucleotides were used according VOGEL et al. (2009): 2.5 μM (forward 5’ ATT ACC GGCGGT TGA GAG G 3’ and 2.5 μM reverse 5’-’ CTG GAA GGA GCT GAG GAA GA 3’ (GenBank Accession Number D49559.3). The PCR conditions were denaturation at 95°C for 5 min, 35 cycles of 95°C for 30 s, 62°C for 30 s, 72°C for 30 s and a final extension at 72°C for 10 min. PCR products were digested for 1U of BssSI restriction endonuclease (Biolabs, New Ipswich, NH, New England) at 37°C for 3h. The genotypes were scored according to restriction fragments length polymorphism (RFLP) analysis as homozygous wild type (150 bp and 95 bp), heterozygous (245 bp, 150 bp, and 95 bp), and mutant homozygous (245bp). Both PCR products and enzyme digestion products were confirmed through electrophoresis on polyacrylamide gels (10%) stained with AgNO_3_. Negative controls (absence of DNA) were applied to ensure that no contaminants were introduced into the reaction.

### Study design and statistical analysis

Anthropometric measurements, sociodemographic information, clinical diagnoses and family history were obtained through an epidemiological questionnaire that had been formulated, tested and applied in our previous study.

The sampling process was random, seeking criteria that would serve a reliable representation of the population of this region. For this study, 4 mL of peripheral venous blood was collected by vacuum method in tubes with ethylenediaminetetraacetic acid (EDTA). A descriptive observational study was performed for the prevalence of the *GIPR* Glu354Gln genetic variant. A control-case study was performed between the cardiometabolic group (n = 146) and control group (n = 156) and the arterial hypertension (AH) group with presence of the C allele (n = 39) and non-carrier AH (n = 66) (Fig 1).

**Fig 1.**
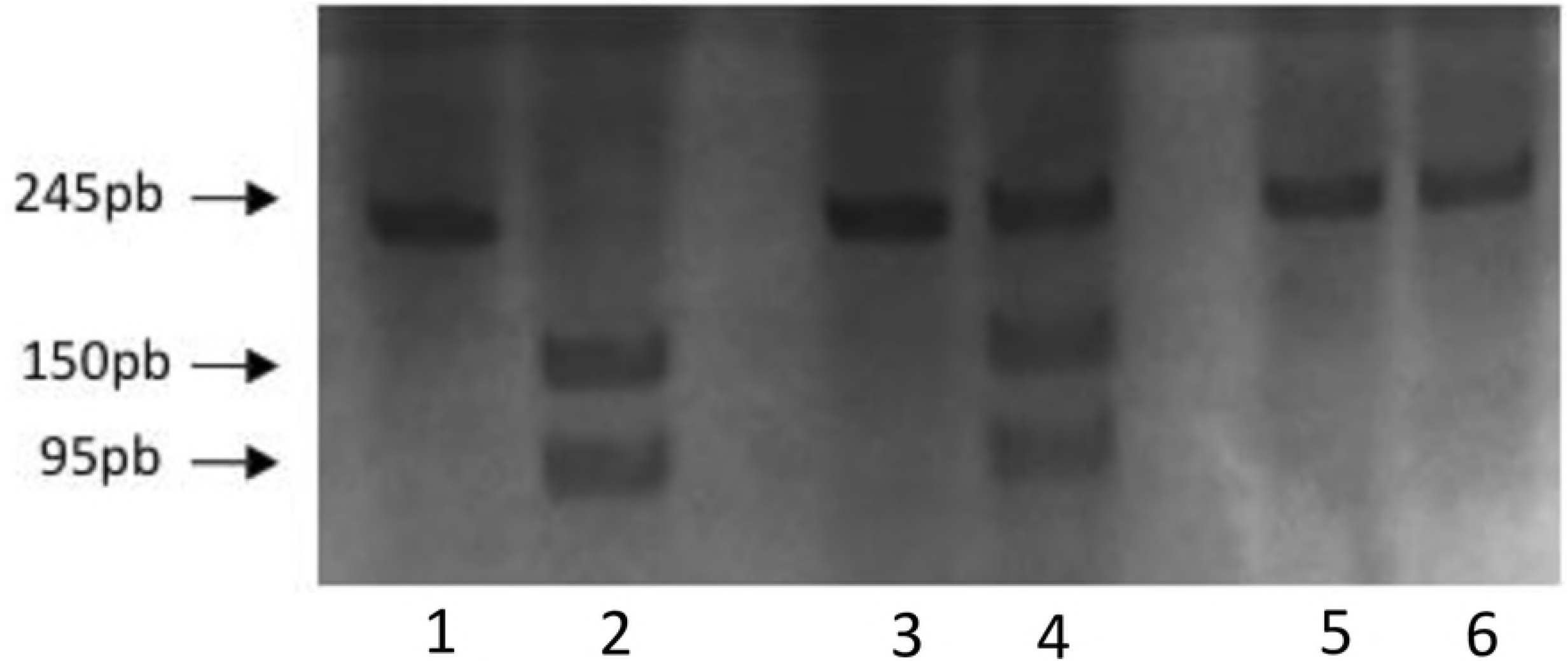
Design of the *GIPR* Glu354Gln polymorphism prevalence study. DCM: cardiometabolic diseases, AH: arterial hypertension.

GraphPad Prism 5.00 (GraphPad Prism Software Inc., La Jolla, CA, USA) and STATA (Corporation Stata Statistical Software Release 13.0. College Station, TEXA, USA) were used to analyze all dataset. Continuous variables were presented as means more or less the standard deviation (SD) and categorical variables as counts (n) and percentages. Comparisons between groups were assessed Student *t*-test, Pearson’s chi-square test and Fisher’s exact test for continuous variables.

## Results

In this study 311 Brazilian participants were selected. The mean ages were 35.1 y (± 16,1) for woman and 46,9 y (± 20,8) for men. Women represented 51,4% (n=160). Sample consisted in 16.6% of individuals from each selected region of the city, being caucasian (67.8%), female (51.5%), adults (61.9%), normal weight (45.0%), sedentary lifestyle (62.1%) and smokers (19.3%). We analyzed the prevalence of the genetic variant of the *GIPR* Glu354Gln, not yet described in this population. Table 1 shows the distribution of genotypes for *GIPR* Glu354Gln polymorphisms, homozygous CC 3,2% (n=10), heterozygous GC 22,5% (n=70) and homozygous wild type GG 74,3% (n=231). The polymorphic C allele carrier was found in 25.7% (n=80) study population. The electrophoretic profile for the genotypes is shown in Fig 2. The samples were tested for Hardy-Weinberg equilibrium, and no deviation from the expected frequencies of genotypes was found for any polymorphism (p>0.05).

**Fig 2.**
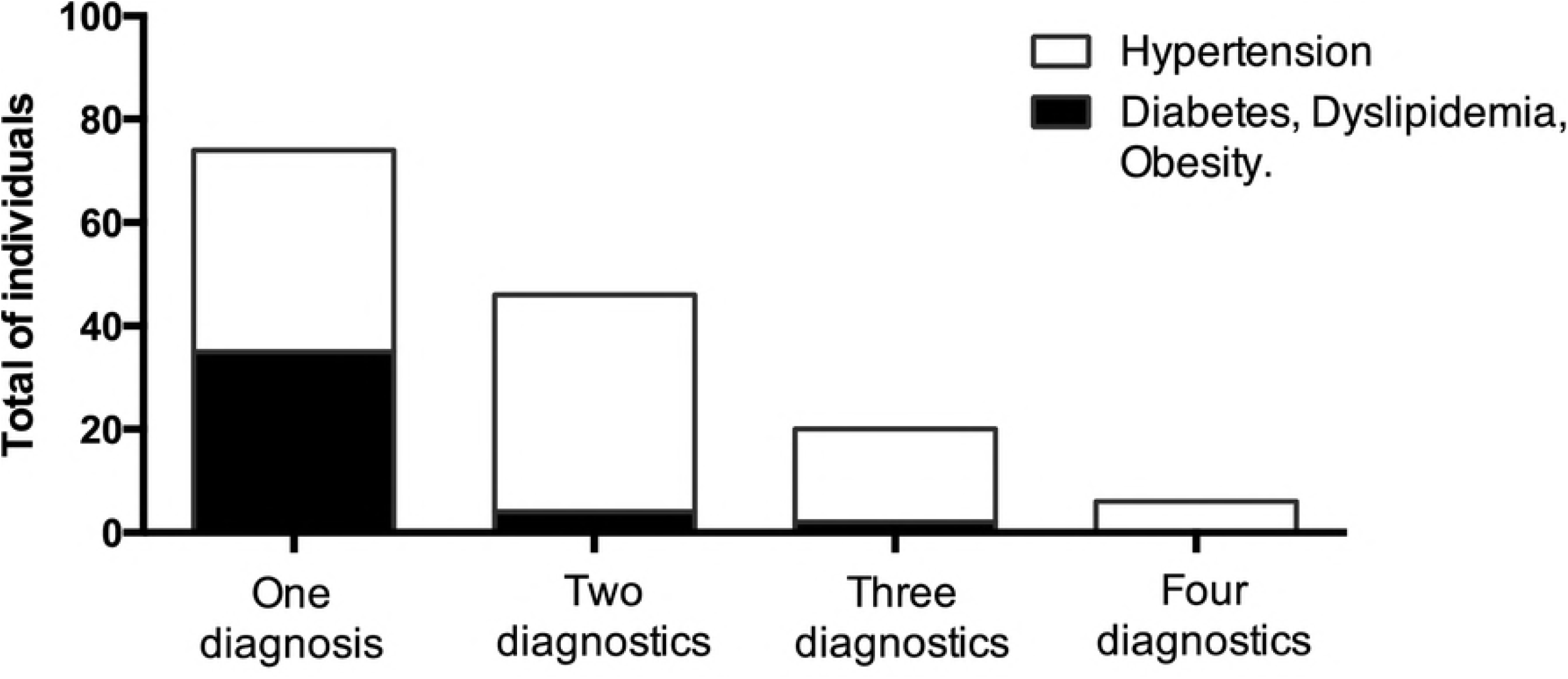
Electrophoretic profile of *GIPR* Glu354Gln gene polymorphism. Genotypes for *GIPR* Glu354Gln: 1, 3, 5: PCR product; 2: GG genotype (150, 95bp); 4: GC heterozygous genotype (245bp, 150bp, 95bp); 6: CC homozygous variant genotype (245bp). 10% polyacrylamide gel electrophoresis, stained with AgNO3.

**Table 1.**
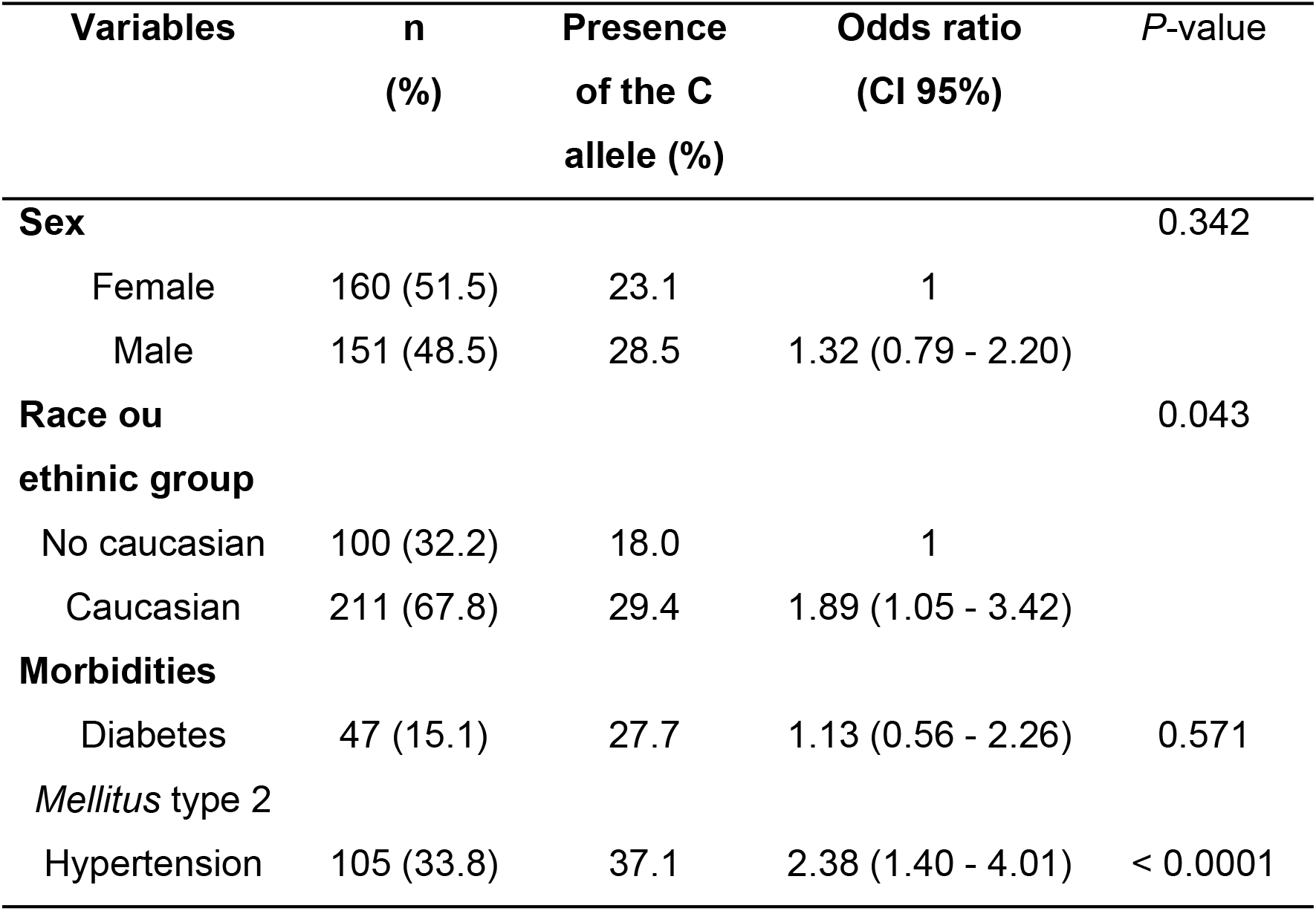
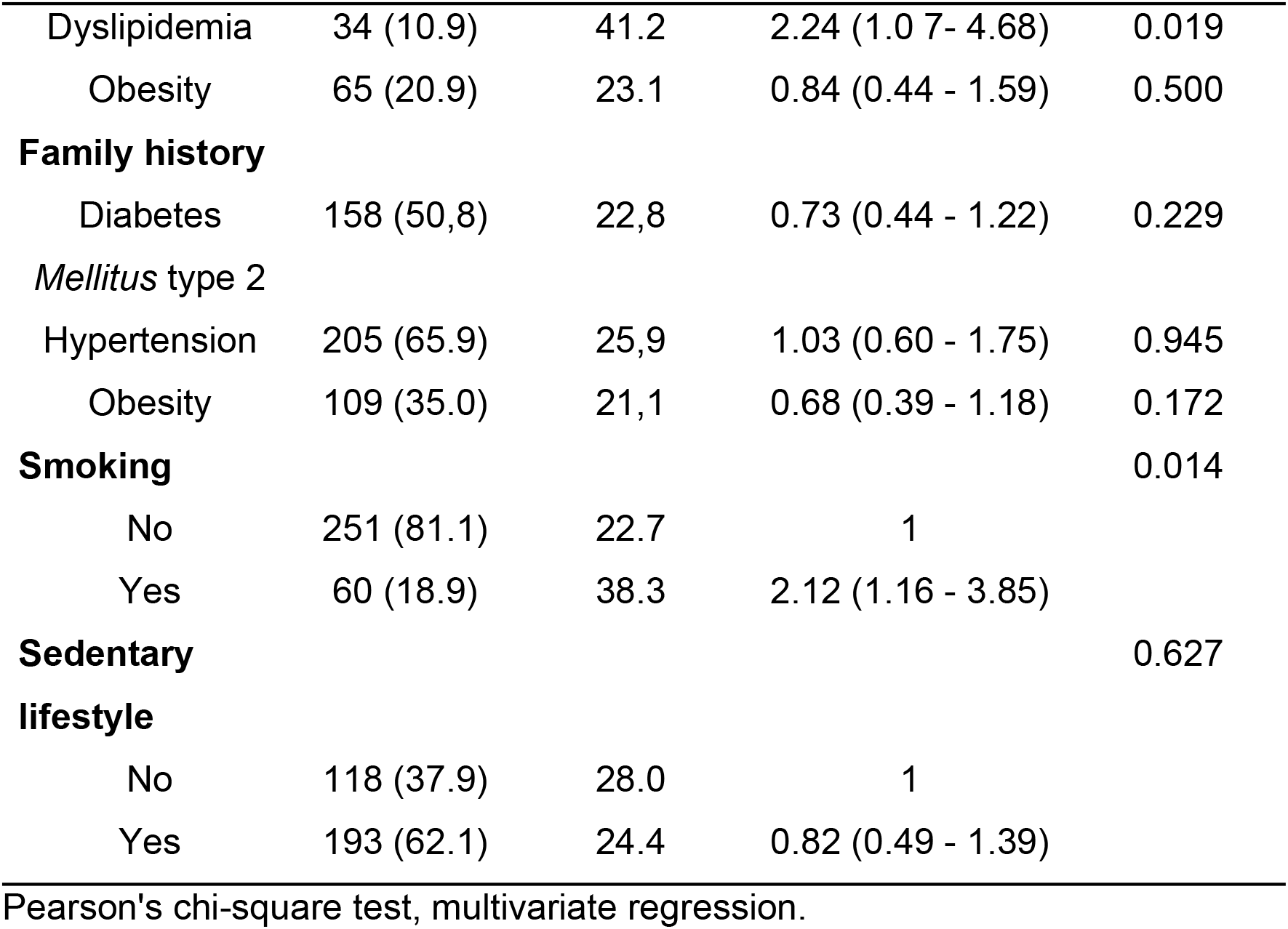
Characterization of the sample for *GIPR* Glu354Gln C allele carriers.

The risk factors recognized for development of cardiovascular disease in this study included diseases or heritability related on glucose metabolism, blood pressure, lipids, weight gain, smoking and physical activity. Hypertension and dyslipidemia showed a significant difference for the for the *GIPR* Glu354Gln variant (Table 1). Highest prevalence rates for the polymorphic allele C were found in the Caucasian 29.4% (*p* = 0.043), hypertensive group 37.1% (*p* < 0.0001), dyslipidemic group 41.2% (*p* = 0.019) and smoking group 38.3% (*p* = 0.014), as showed in Table 2.

**Table 2.** Distribution of the variables according to sex, age and weight status in hypertensive and non-hypertensive individuals

| Variables | Total |  | Women |  | Men |  |
| --- | --- | --- | --- | --- | --- | --- |
|  | AH |  | AH |  | AH |  |
|  | No (n=206) | Yes (n=105)*** | No (n=118) | Yes (n=42)*** | No (n=88) | Yes (n=63)*** |
| **Age, years (mean ± SD)** | 32.33±15.32 | 57.55±15.39**<br>* | 30.03±12.45 | 49.29±16.94*** | 35.42±18,10 | 63.06±11.42* |
| **Body mass index (Kg m<sup>-2</sup>)(mean ± SD)** | 25.12±5.25 | 28.57±6.27** | 25.44±5.89 | 31.38±7.00*<br>** | 24.69±4,22 | 26.70±4.95 |
| **Weight status, n (%)** |  |  |  |  |  |  |
| **Normal weight** | 123 (59.7%) | 36 (34.2%) | 71 (60.2%) | 9 (21.4%) | 52 (59.1%) | 27 (42.9%) |
| **Overweight** | 57 (27.7%) | 45 (42.9%) | 31 (26.3%) | 18 (42.9%) | 26 (29.5%) | 27 (42.9%) |
| **Obese** | 26 (12.6%) | 24 (22.9%) | 16 (13.5%) | 15 (35.7%) | 10 (11.4%) | 9 (14.2%) |

Weight status: normal weight ≥ 18.5 and < 25; overweight: ≥ 25 and < 30; obesity: ≥ 30 (SISVAN, 2004). AH, Arterial Hypertension
Student's t-test, Chi-square test. \*p<0,001, \*\*p=0.0001, \*p<0,0001.

Cardiometabolic diseases were present in 46.9% of the participants (n = 146); arterial hypertension was the most prevalent (33.8%). Hypertension was prevalent in males, advanced age, sedentary lifestyle and smokers. Results indicated that 60% of hypertensive patients (p = 0.004) and 64.7% of dyslipidemic patients (p = 0.046) were male.

There were statistically significant differences in age (*p* > 0.0001) and BMI (p > 0.0001) compared to hypertensive and non-hypertensive subjects, and age was more significant for hypertensive men and BMI than hypertensive women (Table 2).

The family history for diseases (DM2, AH, dyslipidemia and obesity) had no significant association with the *GIPR* Glu354Gln genetic variant. Among participants who presented cardiometabolic diseases (n = 146), AH was the most prevalent disease (71.9%), followed by obesity (43.8%). These clinical conditions were presented alone or in association in which 1.9% of the subjects presented with the four diagnoses. AH was the clinical condition with highest number of associations (Fig 3).

**Fig 3.**
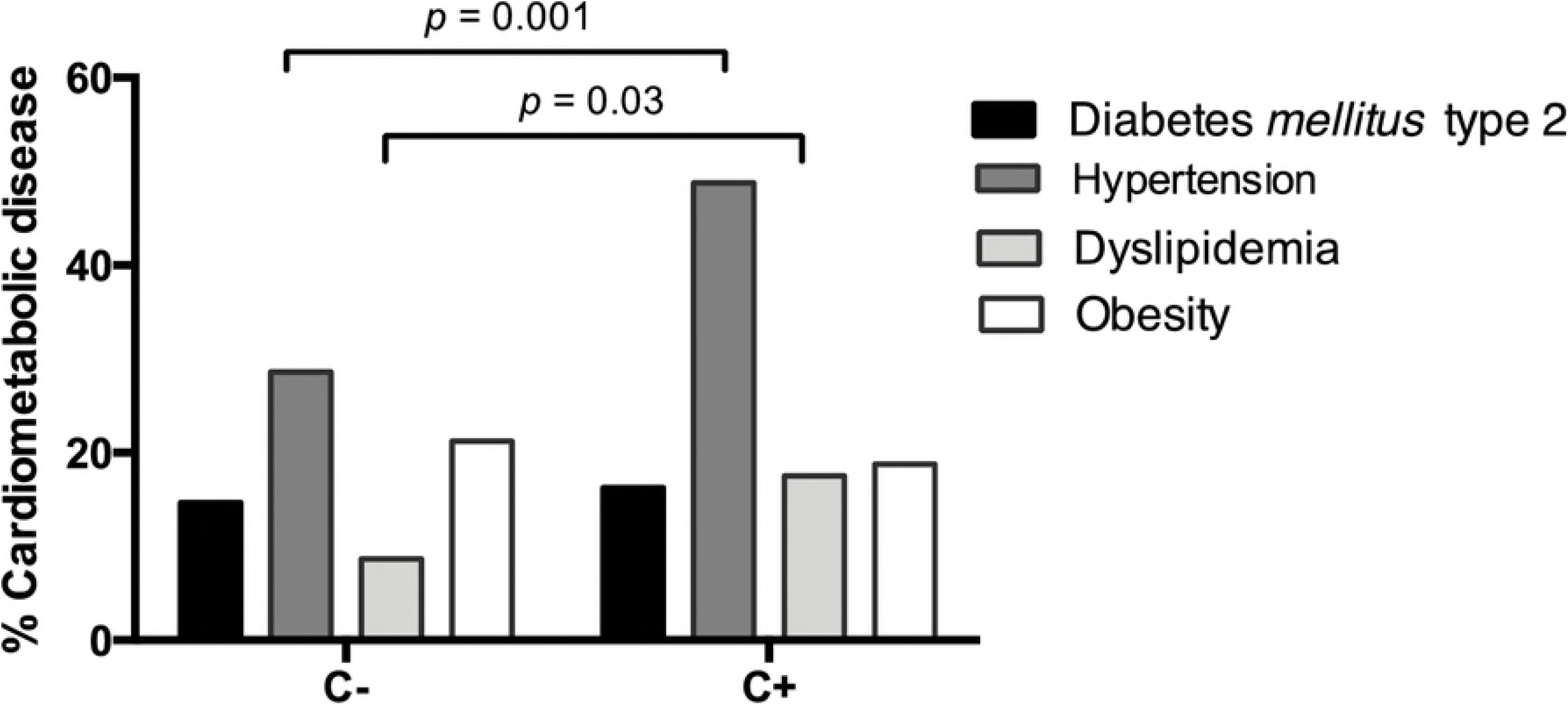
Distribution of diseases according to the number of diagnoses presented. Comparison of AH with the prevalence of other cardiometabolic diseases (n = 146). One diagnosis (individuals with only one disease, without associations), two diagnostics (individuals with two diseases in association), three diagnostics (individuals with three diseases in association in the diagnosis) and four diagnostics (individuals presenting the four associated diseases).

In the next step the case control study was performed. When comparing the control (n = 165) with the cardiometabolic diseases group (n = 146), the genotypic distribution of *GIPR* Glu354Gln showed an associative potential for the dominant model (GC + CC) (*p* = 0.094), supported by allelic frequency (*p* = 0.04) in the cardiometabolic diseases group (Table 3).

**Table 3.** Case-control analyses for *GIPR* Glu354Gln gene polymorphism in the control and cardiometabolic group

| Genetic model | Cardiometabolic diseases |  |  |  |  |  |  |  | p-value |
| --- | --- | --- | --- | --- | --- | --- | --- | --- | --- |
|  | Total |  | NO |  | YES |  | Odds ratio<br>(CI - 95%) |  |  |
|  | n=311 | % | n=165 | % | n=146 | % |  |  |  |
| Co-dominant model |  |  |  |  |  |  |  |  | 0.147 |
| GG | 231 | 74.28 | 129 | 78.18 | 102 | 69.86 | 1 |  |  |
| GC | 70 | 22.50 | 33 | 20.00 | 37 | 25.34 | 0.48(0.11-2.01) |  |  |
| CC | 10 | 3.22 | 3 | 1.82 | 7 | 4.80 | 0.34(0.85-1.34) |  |  |
| Dominant model |  |  |  |  |  |  |  |  | 0.094 |
| GG | 231 | 74.28 | 129 | 78.18 | 102 | 69.86 | 1 |  |  |
| GC+CC | 80 | 25.72 | 36 | 21.82 | 44 | 30.14 | 1.55(0.93-2.58) |  |  |
| Recessive model |  |  |  |  |  |  |  |  | 0.138 |
| GG+GC | 301 | 96.78 | 162 | 98.18 | 139 | 95.2 | 1 |  |  |
| CC | 10 | 3.22 | 3 | 1.82 | 7 | 4.80 | 2.72(0.69-10.71) |  |  |

Cardiometabolic group (n = 146): individuals with diseases related to the cardiometabolic diseases, hypertension, diabetes *mellitus*
type 2, dyslipidemia and obesity. Control group (n = 156).

Regarding *GIPR* Glu354Gln, C allele carriers were twice as likely to present the diagnosis of arterial hypertension and dyslipidemia (Fig 4).

**Fig 4:**
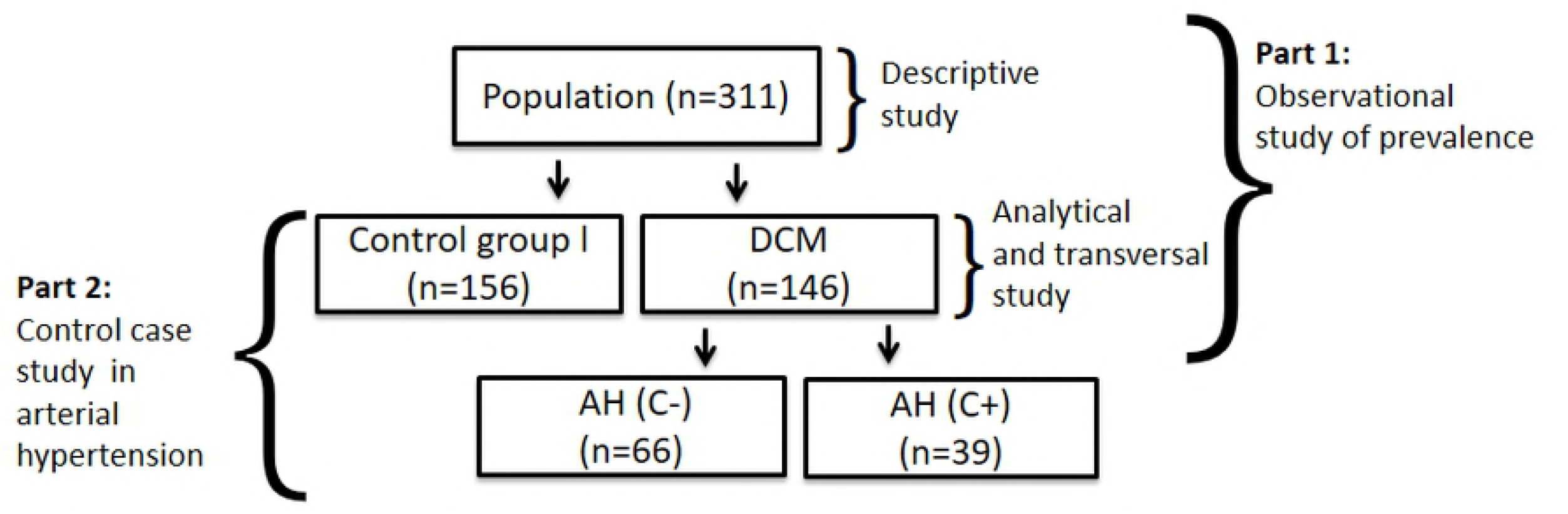
Distribution of clinical conditions related to cardiometabolic diseases in the wild genotype group GG (C-) and the allele C carrier group (C+). No significant association was verified regarding of the *GIPR* Glu354Gln genetic variant and obesity in the total sample population (n = 311) (*p* = 0.639). The distribution of the polymorphic allele between diabetics and non-diabetics did not present statistical difference (*p* = 0,742). Linear regression analysis did not reveal an association between cardiometabolic group and the distribution of genotypes studied, adjusted for age, gender and BMI.

## Discussion

Hypertension is a complex disease that is closely related to endocrine and metabolic disorders (15). The activation of glucose-dependent insulinotropic polypeptide receptor regulates several aspects of pancreatic beta cell and adipocyte metabolism. Some studies have suggested that the functional genetic variant Glu354Gln for the GIPR gene decreases the effect of GIP (8, 10, 12, 16). In the present study involving Brazilian patients with cardiometabolic traits, we demonstrated for the first time that such a genetic GIPR variant has a strong association with hypertension.

In the present work the distribution of genotypes for the GIPR Glu354Gln polymorphism was 3.2% for CC in homozygosis, 22.5% for CG in heterozygosis and 74.3% for GG in homozygous, in a population composed by 67.8% of Caucasian individuals which is a representative sample from inhabitants of the Brazilian State of Paraná according to the national demographic census (17). Regarding the minor C allele, Caucasian individuals presented the highest prevalence rate (29.4 %) in our population. There is no prevalence data for GIPR Glu354Gln in American populations described so far. Few studies presented the GIPR Glu354Gln prevalence in Germany, Netherlands and Denmark (7, 10–12). The polymorphic C allele prevalence in a German population corresponded to 21% of obese children, 20% of obese adults and 22% of the control group (7). In another study with 2280 children and adolescents from Berlin, 6% of obese subjects had a polymorphic genotype, 18% have heterozygous genotype and 61% had a wild genotype (12).

Our study demonstrates the association of the GIPR Glu354Gln polymorphism in the pathophysiology of hypertension and dyslipidemia. The familial history of comorbidities related to cardiometabolic traits (diabetes, hypertension, dyslipidemia and obesity) was not significantly associated with GIPR Glu354Gln genetic polymorphism. Hypertension associated factors were male sex, advanced age, high BMI and smoking, corroborating literature data (18, 19, 20). The mechanisms involved in the pathophysiology of hypertension arise from the interaction among environmental and genetic factors. In a longitudinal study with 6027 Japanese hypertensive patients, PDX1 and LLGL2 genes polymorphisms were associated with HA presence in the dominant model and FAM78B was related to HA in the recessive model. Moreover, the same Japanese study suggests that the presence of BTN2A1 (rs6929846) polymorphic allele predisposes to an inflammatory process acceleration, a genetic risk factor for HA (20). Increased cardiac output in response to higher peripheral vascular resistance occurs with persistent hemodynamic adaptations, even in the absence of clinical manifestations (21).

The lower expression of GIPR in adipose tissue of obese individuals have been related to a decrease in insulin sensitivity resulting in a defect in GIP/GIPR tissue signaling axis (22). Negative correlation was found among GIPR expression in adipose tissue and BMI, abdominal circumference, systolic blood pressure, glycemia and serum triglyceride levels. Our findings did not find an association among the presence of the polymorphism and obesity, however the polymorphic allele was already associated with an increase in insulin resistance in 357 obese children (12).

There is a complex and still poorly understood relationship among GIP and lipid metabolism. The polymorphic allele was more prevalent in individuals with dyslipidemia. It is assumed that this incretin has a vasoactive effect in the adipose tissue, increasing its vascularization, providing greater amount of glucose and esterification of free fatty acids, leading to triglycerides deposition in abdominal subcutaneous tissue (23). In a study using animal models, it was observed that long-term GIP blockade provides an adipose tissue mass reduction and hepatic and muscular triglycerides level reduction, confirming the opposite to expected effect on poor GIP stimulus in this tissue (24). An adverse role of GIP in lipid metabolism was also observed in a Dutch study with diabetic subjects, being that GLP-1, another incretin, would present benefits in lipid normalization of lipids (25). Nitz et al. concluded that there is an LDL levels reduction in heterozygote genotype individuals (11).

Our results showed no difference between diabetic and non-diabetic patients regarding polymorphic allele prevalence, the same result was found in a Dutch study (11).This polymorphic variant was associated with increased peripheral insulin resistance in an obese German children population, suggesting that this polymorphism interferes in glucose metabolism (12). It is known that GIP secretion in response to TTOG and post-diet tests is preserved in diabetics, but with a decrease in GIP effect through a mechanism of downregulation of GIP receptors in pancreatic beta-cells (26). Another GIPR polymorphism, rs10423928, was related to a decrease in insulin response capacity (8).

GIPR Glu354Gln polymorphism showed a threshold association with cardiovascular disease in a Dutch study with 200 individuals (11). This finding suggests that the polymorphic variant has a role in the development of cardiometabolic risk factors. The prevalence of cardiometabolic risk factors in an Iranian study with 5940 adolescents was directly related to a high socioeconomic level (27). Cardiometabolic syndrome was associated with the dominant model, strengthening the hypothesis that the polymorphic variant contributes to the development of type 2 diabetes mellitus, arterial hypertension, dyslipidemia and obesity. We observed the presence of cardiometabolic syndrome in 146 individuals, with hypertension and obesity being the most frequent diseases. Low prevalence of overweight (9.8%) and obesity (1.9%) was observed in a Latin American registry with an adolescent Ecuadorian population, with an important tendency towards glucose intolerance (12.4%) (28). Further studies regarding prevalence of these risk factors in the elderly population are necessary to understand the true impact on the development of cardiovascular diseases and the influence of genetic variants.

## Conclusion

In Brazilian adults, the GG genotype of the *GIPR* Glu354Gln polymorphism was associated with a lower risk of cardiometabolic diseases. This study suggests the involvement of a *GIPR* Glu354Gln in the pathophysiology of hypertension. However, because of low prevalence of the CC genotype, more representative and applicable studies are needed to clarify the mechanisms and underlying genetic effects of our findings.

